# *In vivo* rescue of arboviruses directly from subgenomic DNA fragments

**DOI:** 10.1101/2024.01.17.576038

**Authors:** Maxime Cochin, Jean-Sélim Driouich, Grégory Moureau, Géraldine Piorkowski, Xavier de Lamballerie, Antoine Nougairède

## Abstract

Reverse genetic systems are mainly used to study RNA viruses and rescue recombinant strains in cell culture. Here, we provide proof-of-concept for direct *in vivo* viral generation using the ‘Infectious Subgenomic Amplicons’ method. So far, this procedure allowed to rescue *in vitro* RNA viruses, by the transfection of several overlapping subgenomic DNA fragments encoding the entire virus genome.

We adapted and optimized this technique to generate a pathogenic tick-borne encephalitis virus strain in mice. To define optimal protocol parameters, we injected intramuscularly different amounts of DNA fragments associated, or not, to electroporation. The injection of only 1µg of DNA fragments combined with electroporation resulted in an infection rate of 100%. Then, these parameters were applied to rescue another flavivirus and an alphavirus.

This method provides a novel and efficient strategy for *in vivo* viral generation, which is typically achieved by injecting infectious clones. Furthermore, as part of the development of DNA-launched live attenuated vaccines, this approach, which also has the advantage of not injecting vector DNA, may simplify the generation of attenuated strains *in vivo*.

## Introduction

Development of reverse genetics has greatly facilitated the study of RNA viruses. These tools enable viruses to be recovered and generated by transfecting nucleic acids encoding the entire viral genome into eukaryotic cells.

*In vitro*, numerous reverse genetics methods can be used to rescue RNA viruses from DNA, the most commonly used being the infectious clone method [1]. Briefly, it consists of a single circular DNA molecule containing the complete genome of the studied virus, as well as transcriptional regulatory sequences in the case of RNA viruses. The use of this type of construct requires a bacterial amplification step that can be tedious, if not impossible, due to the toxicity of the viral sequences themselves. To overcome this drawback, numerous methods have been developed since the first “bacteria-free” reverse genetic method defined by Gritsun and Gould in 1995 [2, 3]. Two of these have recently been developed in our laboratory [4, 5]. The ISA method is based on the division of the RNA virus genome into several overlapping subgenomic DNA fragments, flanked by transcriptional regulatory sequences. Subgenomic fragments recovered by PCR amplification are used to generate several RNA viruses by direct transfection into permissive cells. By using several DNA fragments, these methods make it easy to introduce mutations into the viral sequence, which could simplify the development of attenuated strains, potentially for use in vaccinology [6, 7].

*In vivo*, the generation of viruses by the injection of nucleic acids into the cells of living organisms was first studied following the discovery of the infectious power of Tobacco mosaic virus RNA by Gierer and Schramm in 1956 [8]. Subsequently, advances in genetic engineering have enabled the *de novo* generation of viral strains directly in animals from the inoculation of cloned viral genomes within DNA vectors or directly from corresponding RNA transcripts produced *in vitro* [9, 10]. In particular, these techniques have been used to develop live attenuated vaccines (LAV) through the direct injection of DNA [11]. The stability and engineering potential of DNA may make DNA-launched LAV easier to develop and distribute than conventional live attenuated vaccines. Progress in DNA and RNA vaccine approaches has led to the development of numerous tools to improve nucleic acid penetration into targeted cells. For example, chemical reagents such as cationic lipids and polymers are commonly used in the transfection field and physical methods such as electroporation is already employed in clinical trials on DNA vaccines [12–16].

In this work, using the tick-borne encephalitis virus (TBEV, Hypr strain) as a model, we provide proof of concept for the direct *in vivo* application of a recently developed reverse genetics method, the ISA method, by rescuing viral particles after injection of subgenomic amplicons in mice. We also enhance the infection efficiency using an electroporation protocol or a simplified reverse genetic system. Finally, we applied this method to rescue another flavivirus, the japanese encephalitis virus (JEV) and an alphavirus, the chikungunya virus (CHIKV).

## Methods

### Cells

Baby Hamster Kidney 21 cells (BHK21; ATCC-CCL10) were grown at 37 °C with 5% CO2 in minimal essential medium (Life Technologies) supplemented with 5% heat-inactivated fetal calf serum (FCS; Life Technologies), 1% penicillin/streptomycin (PS, 5000 U.mL−1 and 5000 μg mL−1 respectively; Life Technologies), 1% L-Glutamine (Gln; 200 mmol.l−1; Life Technologies) and 1% Tryptose phosphate browth (Trp; Life Technologies).

### ISA method

#### Reverse genetic systems

The reverse genetics systems based on the ISA method were created from the genomes of the JEV_CNS769_Laos_2009 (KC196115), TBEV Hypr (Ref-SKU: 001V-03676; EVAg) and CHIKV LR2006 OPY1 (EU224268). The complete genome of each virus is flanked by the human cytomegalovirus (pCMV) promoter at the 5’ end and by the hepatitis delta ribozyme followed by the simian virus 40 polyadenylation signal (HDR/SV40pA) at the 3’ end. For JEV and TBEV, DNA fragments were synthesized *de novo* (Genscript) and amplified by PCR. For CHIKV, DNA fragments were obtained from an infectious clone. The first fragment was directly amplified by PCR from an infectious clone, which had previously been linearized and digested with the restriction enzyme Dpn1 (New England Biolabs). Purification products of the first CHIKV fragment, which contained a small amount of digested infectious clones, were verified to be non-infectious when tested alone *in vitro*. Second and third fragments were first cloned into a bacterial vector (“StrataClone Vector Mix amp/kan”, Agilent) before being amplified by PCR, as previously described [5].

#### Overlapping DNA fragment preparation

Each DNA fragments was amplified by PCR (Platinum™ SuperFi™ PCR Master Mix and Platinum™ PCR SuperMix High Fidelity; Invitrogene). PCR reactions were performed according to the supplier’s protocol. Briefly, 50µl reactions were conducted with 2µl of DNA template (1ng/µl initial), 2.5µl of forward and reverse primers (10µM initial), 25µl of 2X Mastermix and 18µl of H_2_O in a thermocycler with recommended parameters. Details of primer sequences and PCR cycles are available Table S1. DNA fragments were concentrated and purified by steric exclusion using Millipore 2ml 100KDa columns (Amicon). Briefly, PCR products were placed in the columns and diluted in water. DNA fragments were centrifuged at 3500RPM for 30min, resolubilized in water and centrifuged again at 3500RPM for 30min. DNA fragments were recovered by flipping purification columns and finally centrifuged at 1000G for 1min. Size of DNA fragments were verified using 0.8% agarose gel electrophoresis and their quantity were determined using a nanodrop device. DNA fragments are freeze at −20°C and equimolarly pooled before *in vitro* and *in vivo* experiments.

#### In vitro transfection and viral production

To verify the infective power of the DNA fragments, 96-well plates containing approximately 80% confluent BHK21 cells were transfected with the produced DNA fragments. Briefly, 100ng of equimolar mixture of the produced fragments were transfected using the Lipofectamine 3000 kit (Invitrogen) following the supplier’s recommendations. Viral productions using DNA fragments were performed as previously described and determination of infectious titers was performed as described in the section “Tissue Culture Infectious Dose 50 (TCID50) assay” [5].

### *In vivo* experiments

Experiments were approved by the local ethical committee (C2EA—14) and the French “Ministère de l’Enseignement supérieur et de la Recherche” (APAFIS#9327 and #38899) and performed in accordance with the French national guidelines and the European legislation covering the use of animals for scientific purposes. All animal experiments were conducted in biosafety level (BSL) 3 laboratory.

#### Animal experiments

Three weeks old female C57BL6/J mice were provided by Charles River Laboratories (France). Animals were kept in ISOcage P - Bioexclusion System (Techniplast) with unlimited access to water/food and 14 h/10 h light/dark cycle. Animals were weighed and monitored daily to detect the appearance of any clinical signs of illness/suffering. Virus infection, intramuscular injection and electroporation were realized under general anesthesia (Isoflurane) and organs were collected after euthanasia (cervical dislocation realized under anesthesia).

#### DNA fragments formulation

DNA fragments were formulated in different solutions. NaCl 0.9% was composed of sodium chloride powder and Ultrapure water (Invitrogen). Tyrode’s salt solution was realized using Tyrode’s salts powder (Sigma) and CaCl_2_ (Invitrogene) dissolved in Ultrapure water. Lutrol (Pluronic F-68, Invitrogen) formulation was diluted one day prior inoculation in, NaCl 0.9% or Tyrode salts solution (2X).

#### DNA fragment injection and viral infection

Intramuscular injections of DNA fragments equimolar mixture were performed in posterior tibial muscles of mice (2x50µl).

For electroporation, mice’s right legs were depilated under anesthesia (Isoflurane) one day prior inoculation. On the day of inoculation, 30µl of a hyaluronidase solution (0.4U/µl) diluted in a Tyrode’s salts solution was inoculated at 3 or 4 sites of the tibial muscle. Two to three hours later, a 50µl Tyrode salt solution containing the DNA fragments was inoculated into the same muscle. Immediately after DNA inoculation, a first 95 volts wave, composed of three 100ms pulses separated by 1s interval, was applied on skin surface, around the inoculated muscle, and followed by a second wave applied perpendicularly to the first one. Paracetamol was added to the drinking water for 48 hours after electroporation to reduce animal pain at the injection site. Electroporation was performed with the ECM 830 Square Wave electroporation system and genetrodes (5mm, L-Shape), all provided by BTX group.

For infections using viral productions, animals were inoculated intraperitoneally with doses of 2.10^5^ TCID_50_ for TBEV and 10^4^ TCID_50_ for JEV and CHIKV. Viruses were diluted in a NaCl 0.9% solution.

For JEV and CHIKV, animals were injected with 1mg of anti-IFNAR antibody (Anti-Mouse IFNAR-1 – Purified *in vivo* GOLD™ Functional Grade; Leinco) the day prior and the day following inoculations with infectious material.

A negative control group, named “Mock” was systematically used during animal experiments. These groups were subjected to the same experimental protocol as the experimental groups, except that they were inoculated with a saline solution.

#### Organ collection

Brain of each animal was collected at the time of sacrifice as well as the liver and the spleen of the animals inoculated with DNA or viral form of CHIKV. These organs were then transferred in a 2ml tube containing 1ml of a solution of HBSS 20% SVF and a 3mm tungsten bead. Organs were crushed using a Tissue Lyser machine (Retsch MM400) at 30 cycles/s for 5min and centrifuged at 6000RPM for 10min. The clarified supernatant was transferred into a 1.5 mL tube and centrifuged at 12000RPM for 10 min. Serum was only collected for animals of electroporation studies.

#### Quantitative real-time PCR (RT-qPCR)

The presence of virus in organs of sick animals was confirmed by RT-qPCR. Samples were extracted using the QIAamp 96 DNA kit and RNase-Free DNase Set on the automated QIAcube device (both from Qiagen), following the manufacturer’s instructions. Before extraction, 100μl of each sample was inactivated in a S-Block (Qiagen) loaded with VXL lysis buffer containing RNA carrier and proteinase K. As extraction control, all samples were spiked with 10 μL of internal control (bacteriophage MS2) before acid nucleic extraction as previously described [17].

Real-time RT-qPCR assays (GoTaq 1-step qRT-PCR, Promega) were performed using standard fast cycling parameters (i.e. 10 min at 50 °C, 2 min at 95 °C, and 40 amplification cycles at 95 °C for 3 sec followed by 30 sec at 60 °C) and mix were composed of 3.8 μl of extracted RNA, 6.2 μl of RT-qPCR mix. RT-qPCR reactions were performed on QuantStudio 12 K Flex Real-Time PCR System (Applied Biosystems) and analyzed with the QuantStudio 12 K Flex Applied Biosystems software v1.2.3. Details of detection system are presented in Table S2.

#### Virus isolation assay

The absence of infectious virus in samples from healthy animals was verified by virus isolation assay. Briefly, 96-well plates of confluent BHK21 cells were inoculated with a 5-fold dilution of organ clarified supernatants in culture medium (0% FCS; 1% SP; 1% Gln; 1% Trp) for 2h at 37°C and 5% CO2. The inoculum was then replaced with medium containing FCS (2% FCS; 1%PS; 1% Gln; 1% Trp) and incubated 5 days. Two additional blind passages were conducted using the same protocol. The presence of a cytopathic effects (CPE) was checked at the 3rd passage and the cell supernatants were analyzed by RT-qPCR.

#### Tissue Culture Infectious Dose 50 (TCID50) assay

Infectious viral yields in sick animal organs were assessed by cell culture titration. Infectious titers were determined by inoculating confluent 96-well plates of BHK21 cells with 10-fold serial dilutions of clarified organ supernatants. Each sample was first diluted 100-fold and then serially diluted 10-fold in a 2% FCS medium. After incubation at 37°C and 5% CO2 for 5 days, the presence of a CPE was observed and infectious titers were estimated as TCID50/ml or TCID50/g according to the method of Reed and Muench.

#### Next generation sequencing (NGS) of the full-length genome

Brain supernatant extracts were analysed by NGS and viral subpopulations up to 2% were assessed. Viral RNA was retrotranscribed and amplified using the SuperScript™ III One-Step RT-PCR System with the Platinum™ Taq High Fidelity DNA Polymerase Kit and according to the supplier’s recommendations. Details of the primers used and the amplification cycle are given in Table S3. The PCR products were then purified using the Monarch® PCR & DNA Cleanup Kit (5 μg) according to the supplier’s recommendations. Size of PCR products was checked by agarose gel electrophoresis before sent to the laboratory sequencing platform.

After Qubit quantification using Qubit® dsDNA HS Assay Kit and Qubit 2.0 fluorometer (Thermo Fisher) amplicons were sonicated (Bioruptor®, Diagenode, Liège, Belgium) into 250pb long fragments. Libraries were built adding to fragmented DNA barcode for sample identification and primers with Ion Plus Fragment Library Kit using AB Library Builder System (Thermo Fisher). To pool equimolarly the barcoded samples, a real time PCR quantification step was performed using Ion Library TaqMan™ Quantitation Kit (Thermo Fisher). Next steps included an emulsion PCR of the pools and loading on 520 chips performed using the automated Ion Chef instrument (Thermo Fisher), followed by sequencing using the S5 Ion torrent technology (Thermo Fisher), following manufacturer’s instructions. Consensus sequence was obtained after trimming of reads (reads with quality score <0.99, and length <100pb were removed and the 30 first and 30 last nucleotides were removed from the reads) and mapping of the reads on a reference (GenBank strain: U39292) using CLC genomics workbench software v.21.0.5 (Qiagen). Parameters for reference-based assembly consisted of match score = 1, mismatch cost = 2, length fraction = 0.5, similarity fraction = 0.8, insertion cost = 3, and deletion cost = 3. A de novo contig was also produced to ensure that the consensus sequence was not affected by the reference sequence. Quasi species with frequency over 2% were studied.

#### In vitro replication fitness

96-well plates of confluent BHK21 cells were infected with clarified brain supernatants at an MOI of 0.01. Cells were infected with a 20µl inoculum for 2 hours. The inoculum was removed, cells were rinsed with HBSS and then 150µl of 2% FCS medium was added to each well. Replication kinetics were performed in triplicates for each sample and for a period of 3 days. Each collection was performed on independent wells. Each day, 100µl of culture supernatant was collected and analyzed by RT-qPCR as described in the section “Quantitative real-time RT-PCR (RT-qPCR)”. The detection and quantification system used is available in Table S4. RNA quantities were determined using four serial dilutions of an appropriate quantified T7-generated synthetic RNA standards.

#### Graphical representations and statistical analysis

Graphical representations and statistical analyses were performed with Graphpad Prism 9 (Graphpad software) except the calcul of 50% lethal dose (LD50) of DNA fragment that was performed using the AAT bioquest online tool (www.aatbio.com/tools/ld50-calculator). Statistical details for each experiment are described in corresponding Supplementary tables. Survival curves comparison were obtained by Kaplan-Meier analysis and viral RNA yields were analysed using a Two-way ANOVA tukey’s multiple comparisons test. P-values lower than 0.05 were considered statistically significant. Experimental workflow and reverse genetic systems schemes were created on biorender.com.

## Results

### Proof of concept of the *in vivo* ISA method

The efficacy of the *in vivo* application of the ISA method was evaluated by injecting overlapping subgenomic DNA fragments, encoding the full-length genome of TBEV, directly into mice (Figure 1). Each subgenomic DNA fragment was *de novo* synthetised and amplified by PCR on a separate plasmid. The full-length DNA genome is flanked by a eukaryotic transcription promoter at the 5’ end and by the hepatitis delta ribozyme followed by the simian virus 40 polyadenylation signal (HDR/SV40pA) at the 3’ end.

**Figure 1:**
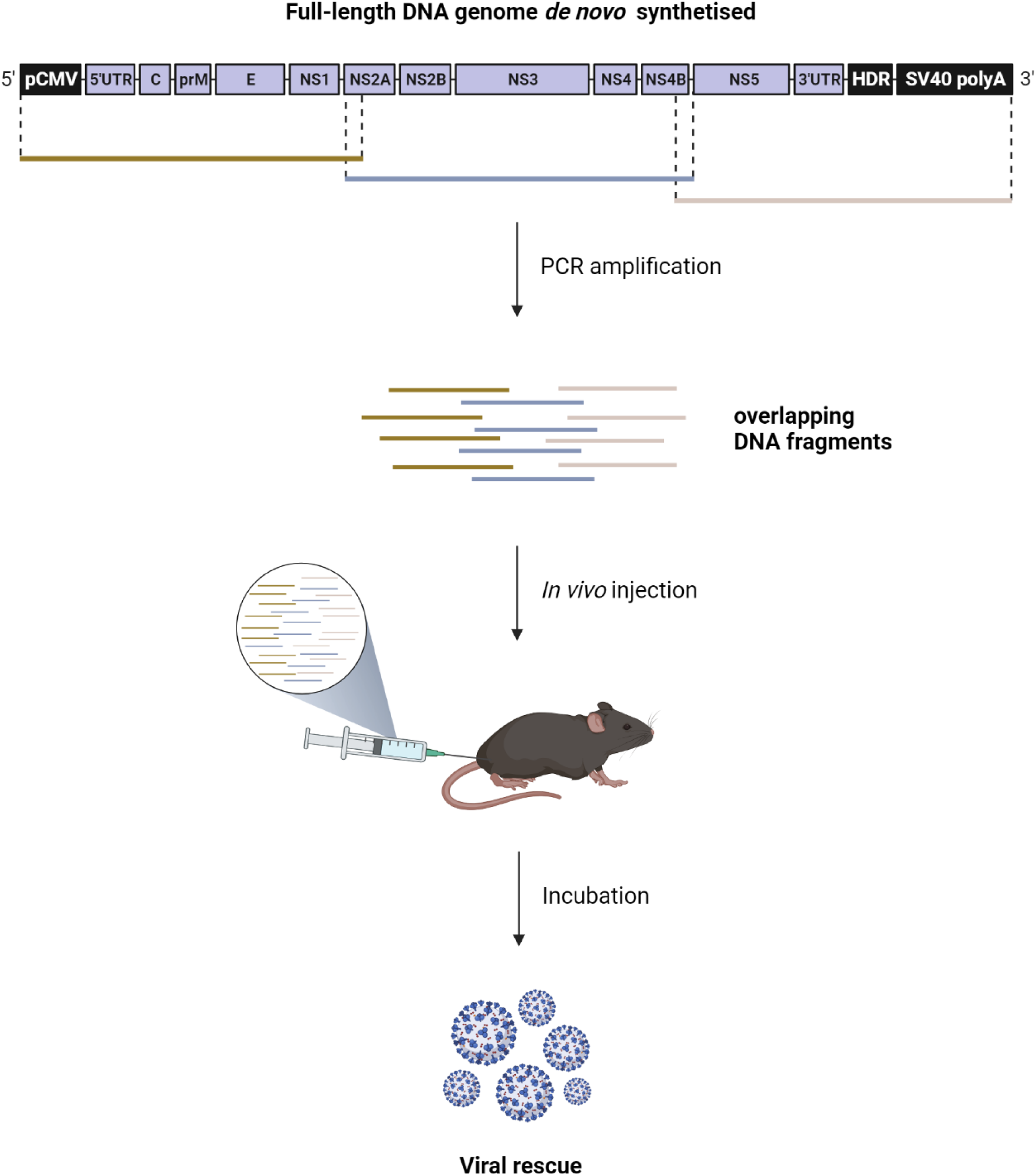
*In vivo* ISA method: The subgenomic DNA fragments amplified are equimolarly pooled and directly injected into animals. After an incubation period, animals show signs of viral infection that coincide with virus detection.

For this proof of concept, we used the Hypr strain of the TBEV (Figure S1). Groups of 6 mice were inoculated in the posterior tibial muscle with 3 subgenomic DNA fragments, at different doses, formulated in NaCl 0.9% solution. The mice were monitored daily for 21 days for signs of infection. Survival curves were obtained on the basis of suffering criteria. The combined results of two independent experiments are shown in figure 2.

**Figure 2:**
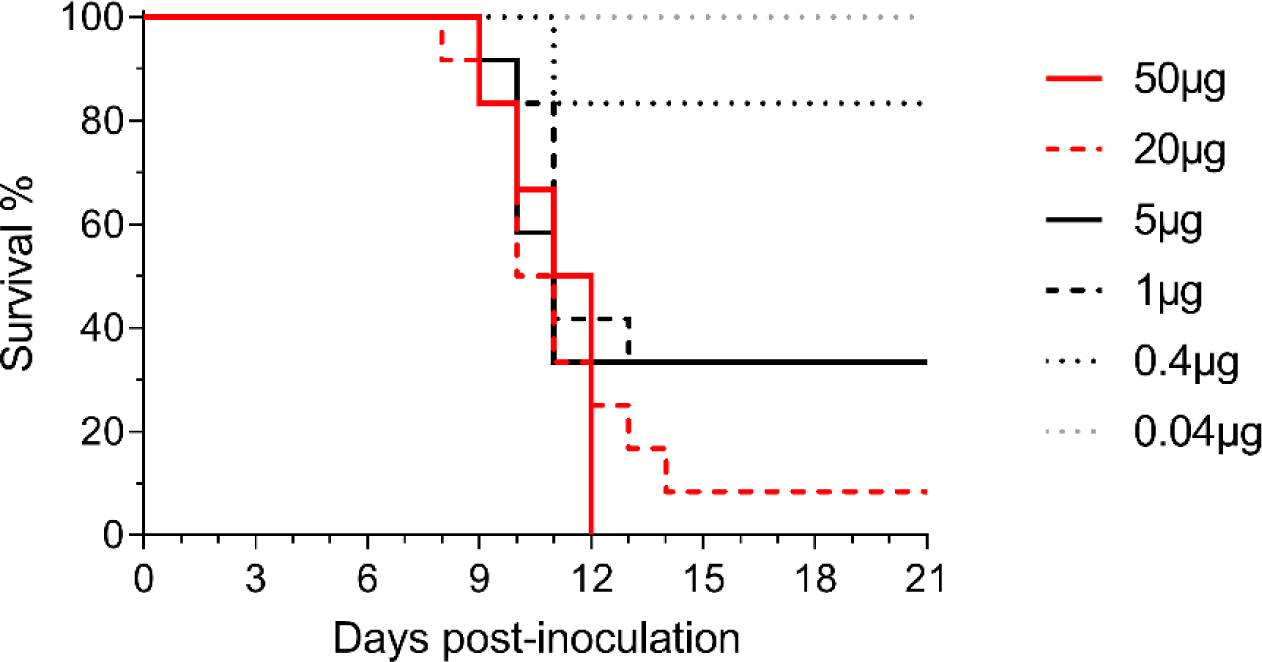
Survival curves after the inoculation of TBEV DNA fragments *in vivo*. Survival data are presented as Kaplan-Meier curves. Survival curves represent results from six different doses of DNA fragments formulated in NaCl 0.9%: 50, 0.4 and 0.04µg (n=6 animals) and 20, 5 and 1µg (n=2x6 animals). Statistical analysis are presented in Table S5. Related infectious titers in brains for these groups are presented in Figure S2.

Results showed that the *in vivo* ISA method is effective: inoculation of mice with DNA fragments resulted signs of viral infection, in all groups, except the 0.04µg dose. At a dose greater than or equal to 1 µg, more than half the animals were infected. The rate of infection is proportional to the quantity of DNA fragments inoculated (Figure 2). These observations allow us to establish a 50% lethal dose (LD50) of 0.67µg. The presence of infectious virus in the brains of sick animals was confirmed by cell culture titration (Figure S2). The absence of virus in the brains of surviving animals was confirmed by viral isolation assays at the end of follow-up.

In order to optimize the rate of infection in our model, other formulations were tested. The use of solutions of Tyrode’s salts or Lutrol 3.5mM, a cationic polymer, did not improve the rate of infection (Figure S3).

We previously demonstrated that the fidelity of the PCR polymerase used to produce subgenomic DNA fragments influences the genetic variability of the viral populations rescued *in vitro* [18]. To investigate this question *in vivo*, we inoculated mice with 20 µg of DNA fragments amplified using a high-fidelity polymerase (Taq HiFi, 6 times fidelity of Taq polymerase) or a super high-fidelity polymerase (Taq SuperFi, over 300 times fidelity of taq polymerase). We sequenced the complete genomes of viruses present in the brains of sick mice and did not observe major differences between groups. (Figure S4. A-B). We also evaluated the in vitro replicative fitness of viruses collected from the brain of one animal in each group and did not observe important differences once again. (Figure S4. C). Overall, there are no major differences in genomic and fitness characteristics between viral populations generated using a high-fidelity or super high-fidelity polymerase. For the following experiments, we decided to use the super high-fidelity polymerase.

### Optimization of the *in vivo* ISA method: electroporation and reduction in the number of DNA fragments

To improve viral production using the *in vivo* ISA method, we explored two potential optimization strategies: (1) coupling intramuscular injection with an electroporation protocol, and (2) using two overlapping subgenomic DNA fragments (Figure S5).

Groups of 6 to 18 mice were inoculated with amounts of DNA fragments ranging from 1 to 0.02µg. For each experiment, a control group of 6 mice received 1µg of DNA fragments intramuscularly. As previously, survival curves were obtained on the basis of suffering criteria and the presence of infectious virus was confirmed in the brains of sick animals, as well as the absence of virus in survivors (Figure S6 and S7). Based on these data, rate of infections (i.e. % of confirmed infected animals) are summarized in Figure 3.

**Figure 3:**
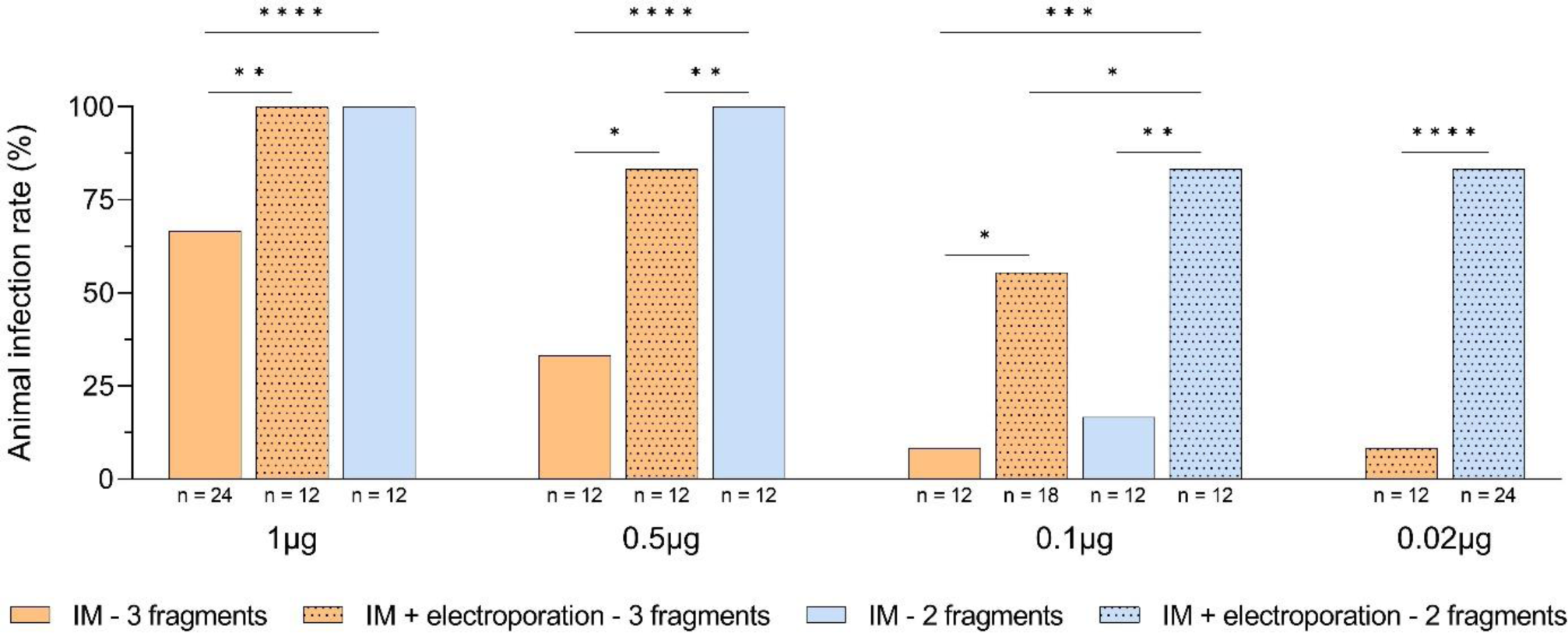
Rate of infections after optimization of the *in vivo* ISA method. Each bar represents the rate of infection (i.e. % of confirmed sick animals) for each conditions explored. Number of mice (n) in each condition is show below the X-axis. This figure shows data presented in Figure S7 and Table S6. Statistical analysis were performed on survival curves in Figure S7 and presented in Table S7. ****, ***, ** and * symbols indicated a significant difference with a p-value ≤0.0001 and ranging between 0.0001–0.001, 0.001–0.01, and 0.01–0.05, respectively.

With equal amounts of DNA fragments, both electroporation protocol and the use of two DNA fragments resulted in a better rate of infection. Rates of infections were 1.5 to 6.7 times higher by using electroporation. This improvement was significant at all doses when using 3 (p=[0.01;0.0059]) and 2 (p=0.0036) fragments. Rates of infections were 1.5 to 10 times higher when using two DNA fragments instead of three. This improvement was significant at 1µg and 0.5µg doses (p≤0.0001) when using IM injection and at 0.1µg and 0.02µg doses (p≤0.0325) by using electroporation.

### Application of the *in vivo* ISA method to other viruses

To evaluate the versatility of the *in vivo* ISA method, we used the same approach with two other arboviruses: JEV and CHIKV (Figure S8).

One day after receiving anti-IFNAR antibody, groups of 12 animals were inoculated with 1µg of DNA fragments by intramuscular route coupled with electroporation and control groups of 6 mice were inoculated with a dose of 10^4^ TCID_50_ of JEV or CHIKV produced *in vitro* by the ISA method. Once again, survival curves were obtained in the basis of suffering criteria (Figure 4).

**Figure 4:**
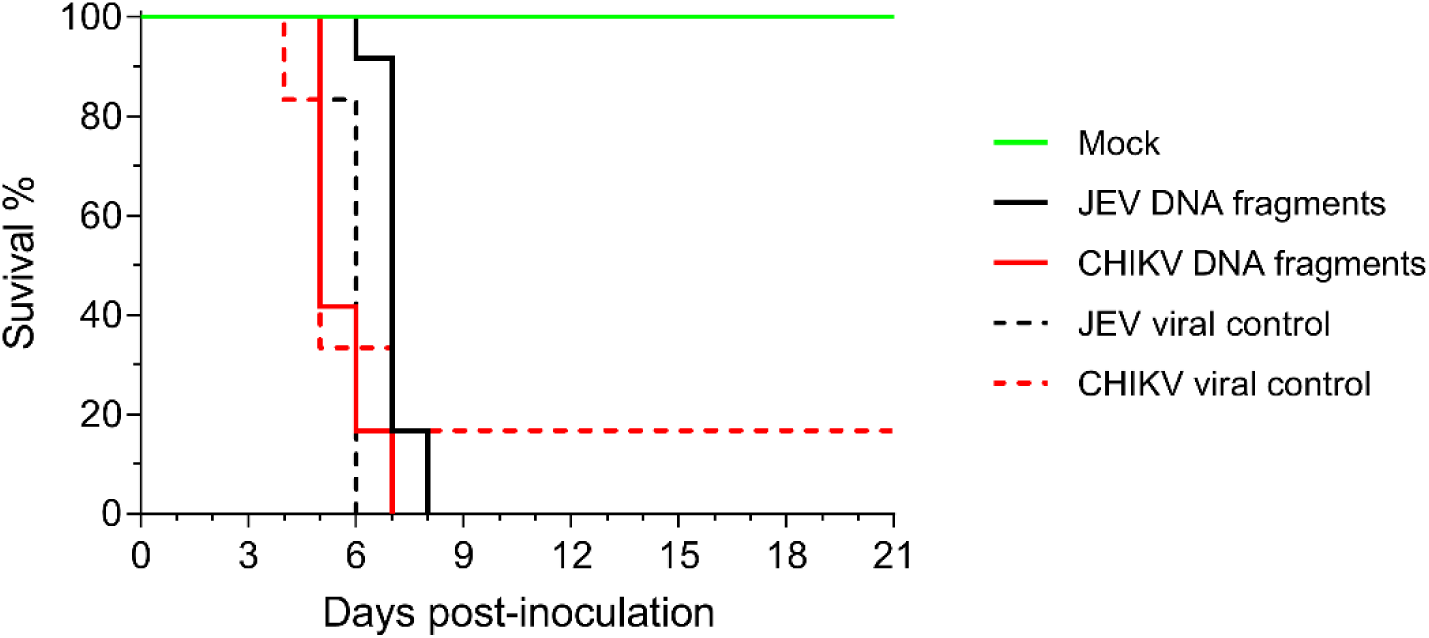
Survival curves for JEV and CHIKV infected animals. Survival curves of animals infected by the inoculation of viral particles produced *in vitro* by the ISA method (dotted lane; n=12) or by the *in vivo* ISA method (lane; n=6).

Results showed that the *in vivo* ISA method is fully applicable to other arboviruses: inoculation of DNA fragments resulted in signs of infection in all groups. Injection of the JEV strain produced *in vitro* caused infection in all animals, while only one survived in the group inoculated with the CHIKV strain. Brains were recovered from all animals. Spleens and livers were also harvested from CHIKV-infected animals. The presence of virus in organs of sick mice was confirmed by RT-qPCR and quantified by cell titration (Figure S9).

## Discussion

The ISA method is a simple procedure for rescuing viruses *in vitro* within days. It involves the direct transfection of susceptible cells with overlapping subgenomic DNA fragments covering a complete viral genome. In this study, we adapted this method to produce the Hypr strain of the TBEV directly *in vivo* and demonstrated its effectiveness. Fragments of subgenomic DNA, identical to those used in vitro, were directly inoculated intramuscularly into mice. This method was effective to generate *de novo* infectious viruses in mice with doses ranging from 50 to 0.4 µg formulated in a 0.9% NaCl solution. We also varied the parameters of this protocol to conclude that the use of an electroporation protocol and a lower number of fragments (2 instead of 3) increased the rate of infection of the animals and achieved 100% efficacy with 1µg doses. By combining these most effective parameters, we were able to rescue two flaviviruses and one alphavirus: TBEV, JEV and CHIKV.

Numerous RNA and DNA viruses have already been produced *in vivo* following inoculation of nucleic acid into animals. However, it generally involves inoculation of the viral genome as a single molecule, an infectious clone in most cases. To our knowledge, two other studies have reported the inoculation of fragmented genetic material to rescue viruses. In the 90’s, Sprengel et al. observed, in a duck model, the generation of infectious duck hepatitis B virus particles, through recombination of DNA fragments [17]. In a more recent work, Willems et al. observed a similar phenomenon with the bovine leukemia virus through inoculation of clones that were individually non-infectious but capable of producing viable viruses when injected simultaneously [19].

The main current application of DNA-launched *in vivo* infection is the development of new strategy to deliver LAV. Several studies have shown that DNA coding for live attenuated viral strains of several arboviruses, such as yellow fever virus and venezuelan equine encephalitis virus, can induce a protective immune response in mice [20–32]. Some of these viruses were also generate *in vivo* using an electroporation protocol following intramuscular inoculation of DNA. However, these studies use an infectious clone as genetic material for transfection, which has certain disadvantages for their production within a bacterial vector.

The ISA method has the capacity to generate viral strains quickly and easily *in vitro*. The use of several subgenomic DNA fragments makes it possible to introduce mutations into the genome of the virus under study, thus producing attenuated viral strains. We believe that the *in vivo* transposition of this technique, presented in this study, will provide the same advantages. As well as the ability to directly assess the immunogenicity of these attenuated viral strains, making this a new tool for the development of LAV.

With the exception of transcriptional regulatory sequences, the genetic material used in the *in vivo* ISA method corresponds solely to the complete genome of the virus of interest while genomic sequences from other prokaryotes and eukaryotes were also administrated when using an infectious clone. Indeed, this is part of the current debate on the injection of foreign genetic material into humans. Nevertheless, this particularity does not prevent this method from being carefully evaluated with a view to its safe use in humans.

## Author contributions

MC, XdL and AN designed the paper. MC, GM, JSD and GP performed and analyzed experiments. MC wrote the paper. AN and JSD reviewed the paper. MC, XdL and AN designed and supervised experimental work. All authors have read and approved the submission of the manuscript.

## Acknowledgements

We thank Rayane Amaral and Noémie Courtin (UVE; Marseille) for them technical contribution. This work was supported by the ANRS-MIE (PRI project of the EMERGEN research program) and by European Virus Archive Global (EVA 213 GLOBAL) funded by the European Union’s Horizon 2020 research and innovation program under grant agreement No. 871029.

## Data availability statement

Authors can confirm that all other relevant data are included in the paper and/or its Supplementary information files.

## Supplementary information

**Figure S1:**
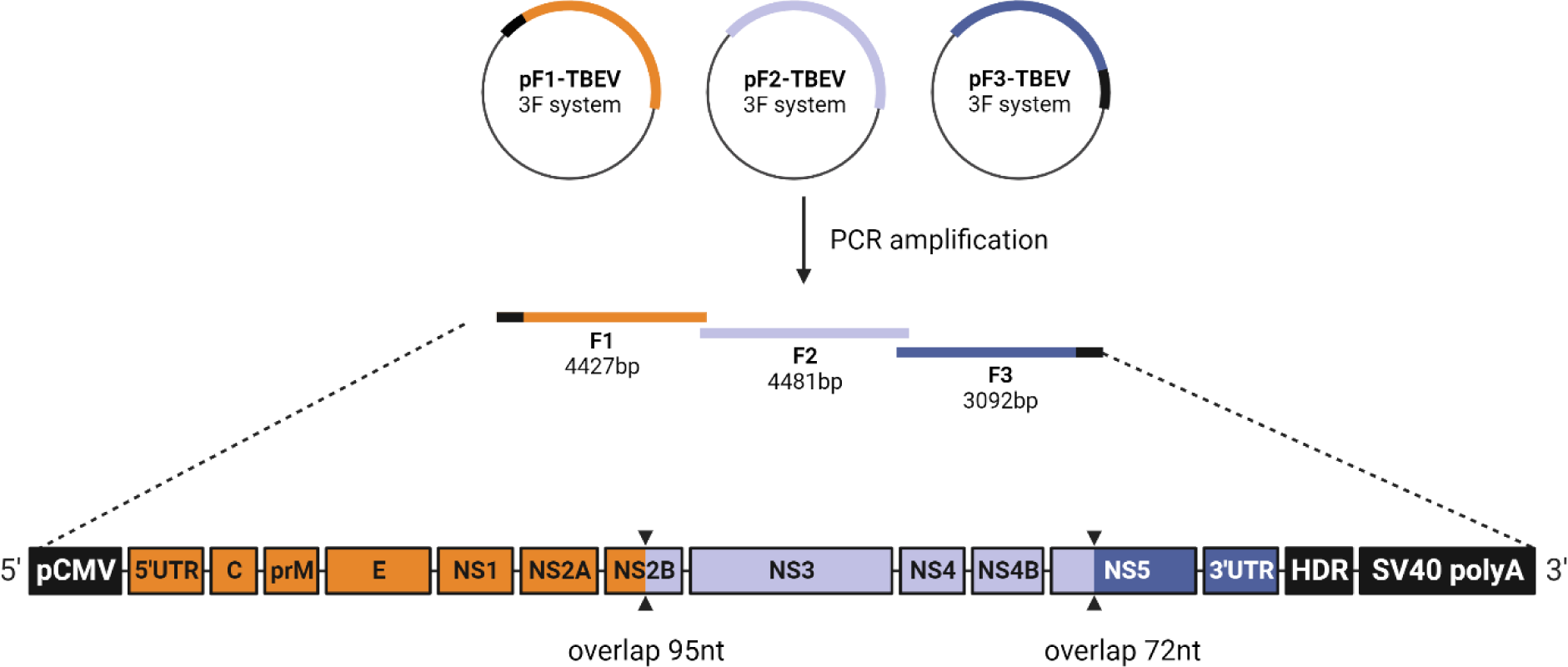
Three-fragmented reverse genetic system used to generate the Hypr strain of TBEV. DNA fragments were *de novo* synthetised and cloned into plasmid (Genscript).

**Figure S2:**
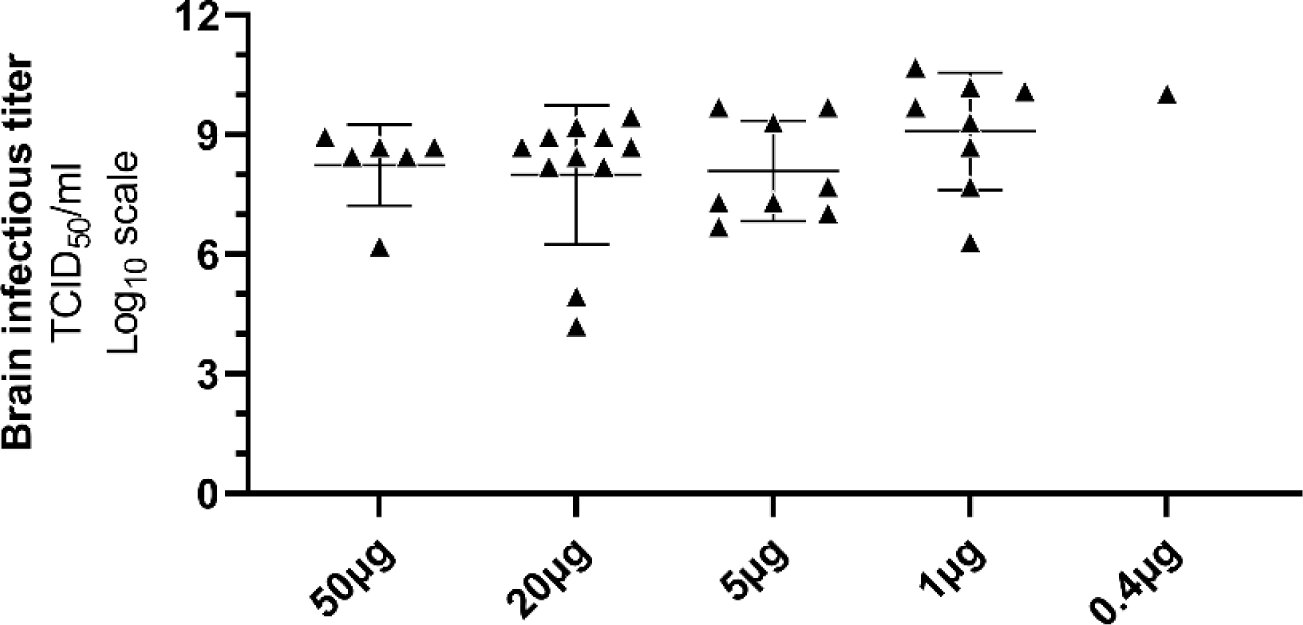
Brain infectious titers of infected animals presented in figure 1.

**Figure S3:**
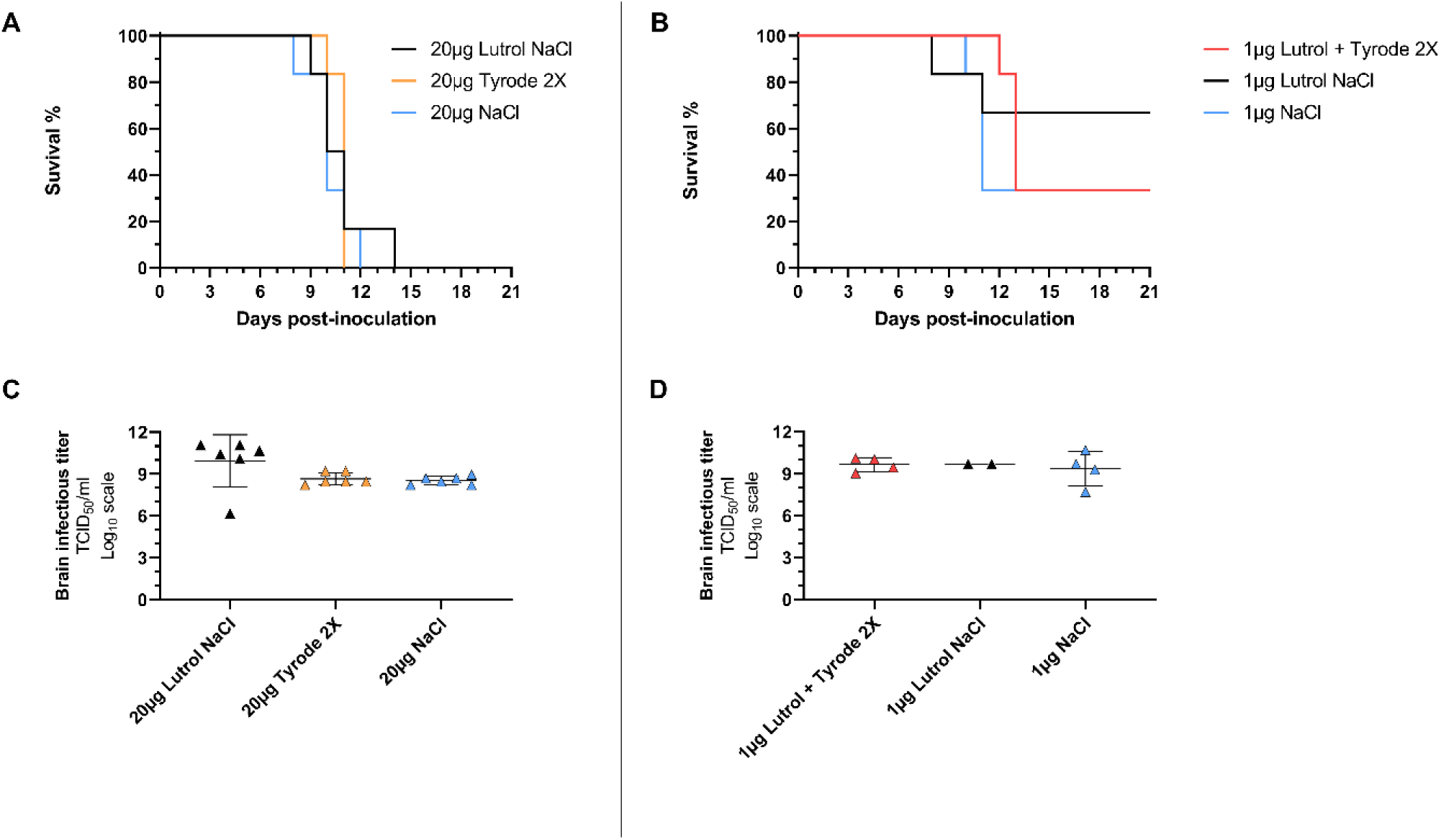
Assessment of different formulations to improve viral rescue. Figure presents data obtained from 2 experiments with groups of 6 mice (n=6): survival curves (A-B) and infectious titers in brain of infected animal (C-D).

**Figure S4:**
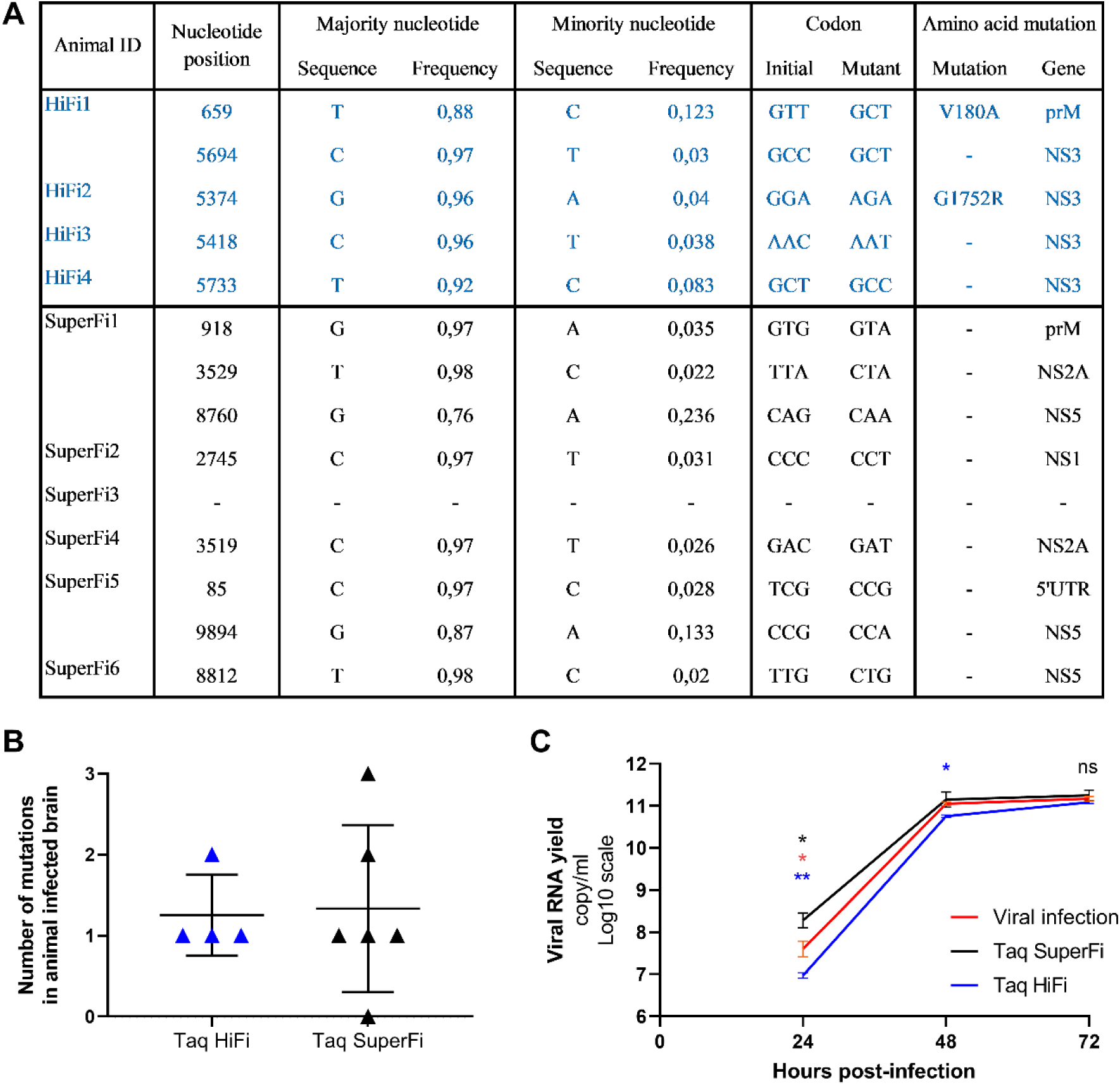
Viral rescue *in vivo* using different polymerase for DNA fragments amplification. Data represent mutational analysis by next-generation sequencing (A, B) and *in vitro* replication fitness (D). *In vitro* replication fitness was conducted in triplicates on one animal sample per condition. As a control, we also tested the replication fitness of a viral population recovered from a clarified brain supernatant intraperitoneally infected with the Hypr strain of TBEV (2.10^5^ TCID_50_). Statistical analysis of replication fitness assay is detailed in Table S8. ** and * symbols indicate a p-value ranging between 0.001–0.01, and 0.01–0.05, respectively. Blue digits correspond to comparison between “Taq SuperFi” and “Taq HiFi” values. Red digits correspond to comparison between “Taq SuperFi” and “Viral infection” values. Black digits correspond to comparison between “Taq HiFi” and “Viral infection” values.

**Figure S5:**
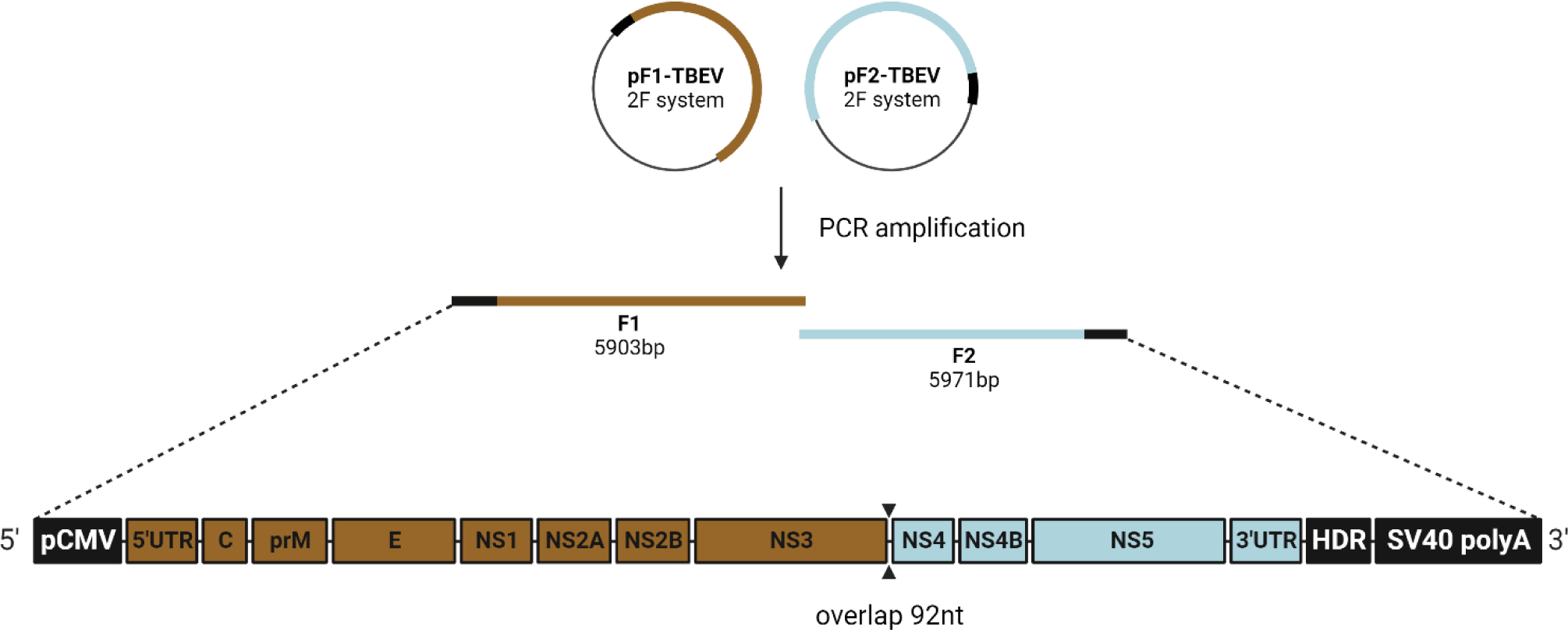
Two-fragmented reverse genetic system used to generate the Hypr strain of TBEV. Genetic sequence is exactly as in the TBEV three-fragmented system.

**Figure S6:**
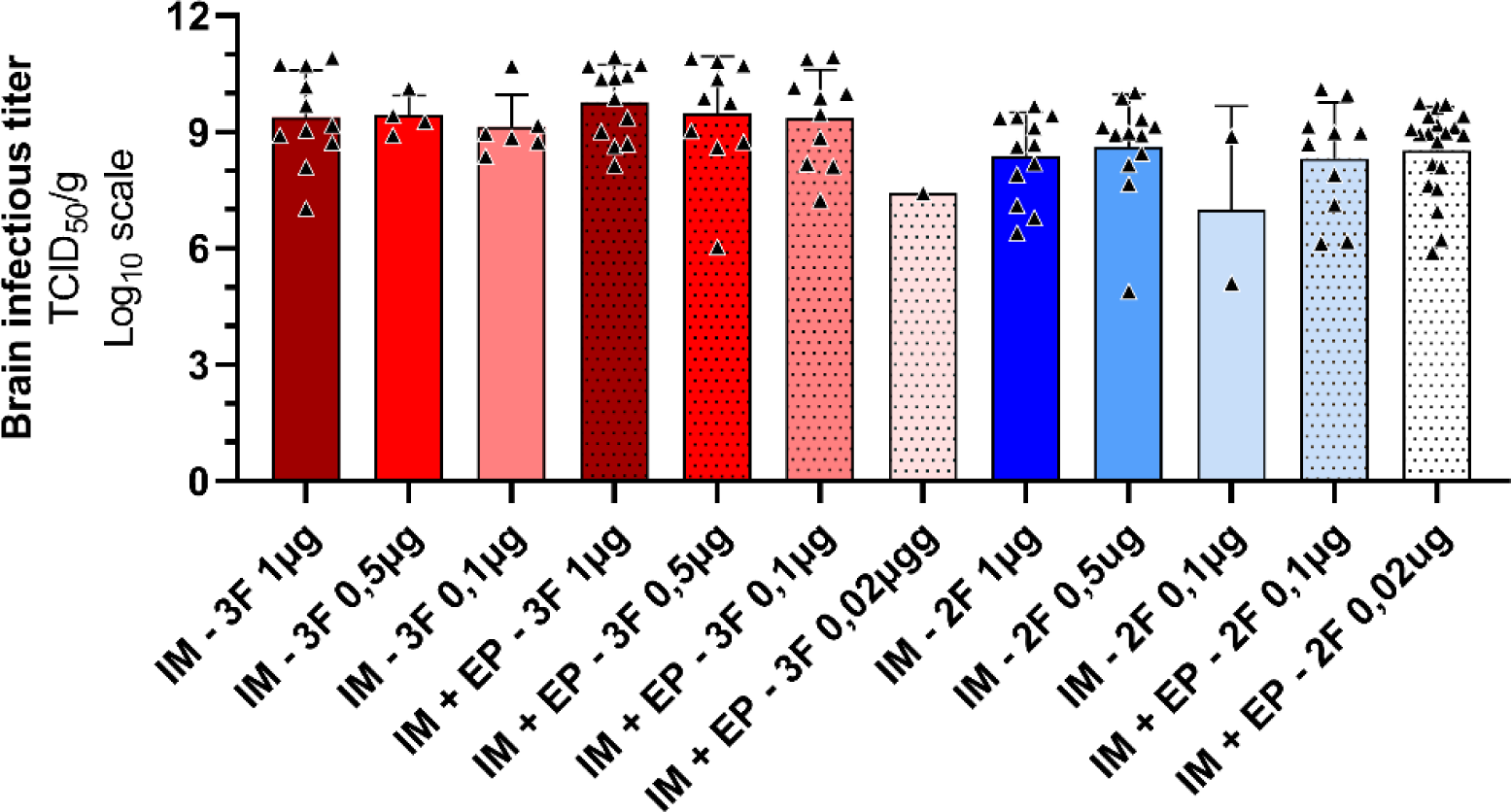
Brain infectious titers of mice infected during experiments presented in figure 3. (IM = Intramuscular; EP = Electroporation; 3F = use of three fragments; 2F = use of two fragments)

**Figure S7:**
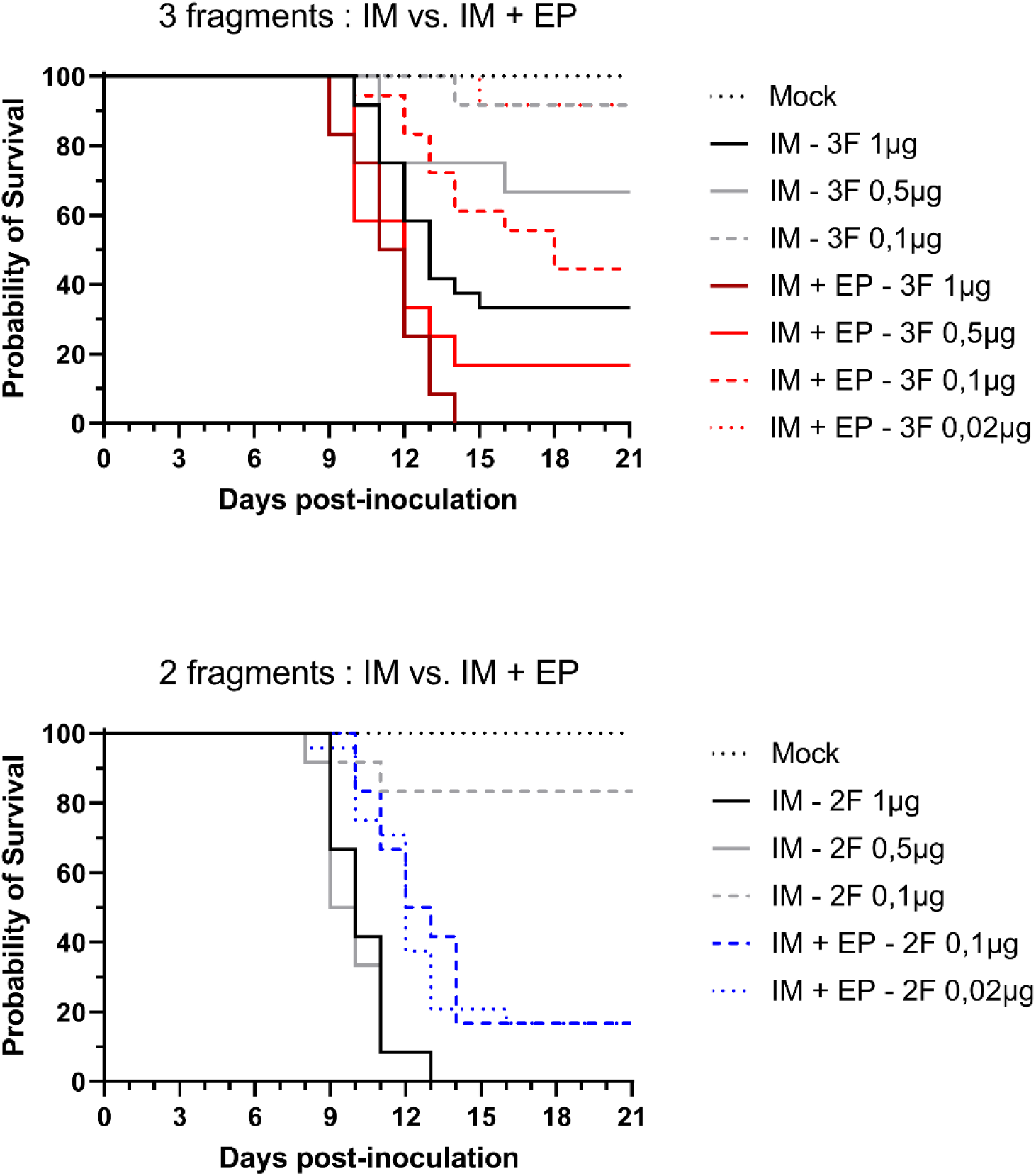
Survival curves used to present results in figure 3. (IM = intramuscular; EP = Electroporation; 3F = three-fragmented; 2F = two-fragmented). Graphs shows data pooled from independent experiments. Statistical analyse are detailed in Table S7.

**Figure S8:**
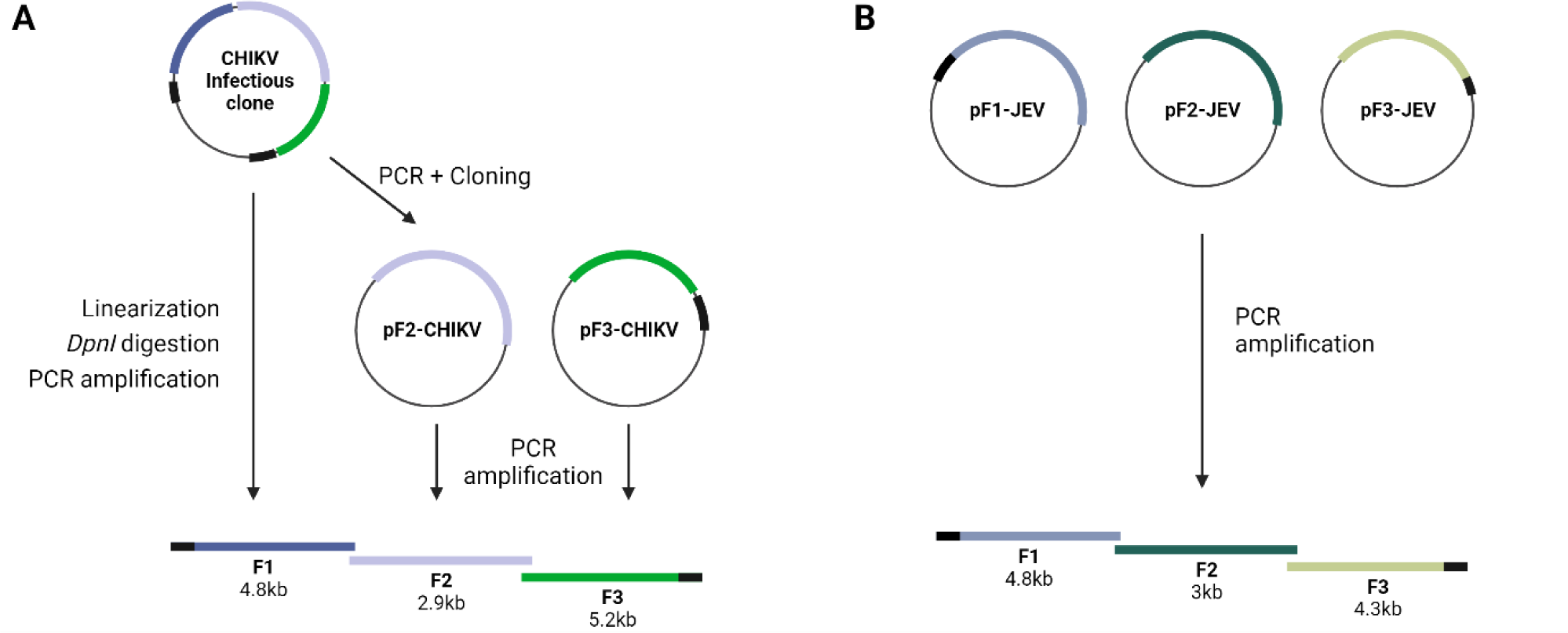
Three-fragmented reverse genetic systems used to generate CHIKV and JEV. CHIKV (A) and JEV (B) reverse genetic systems.

**Figure S9:**
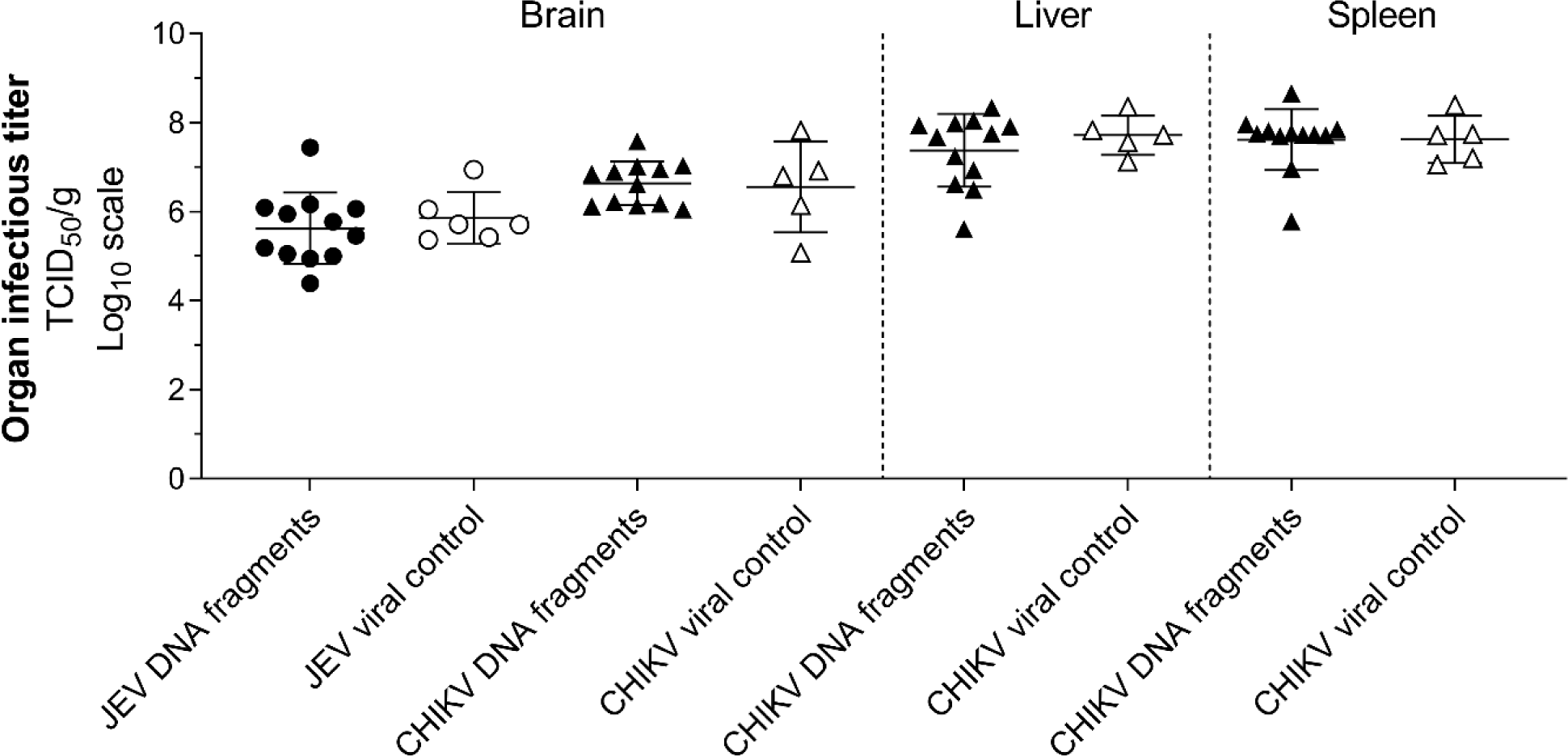
Infectious titers measured in organs of JEV and CHIKV infected animals.

**Table S1:**
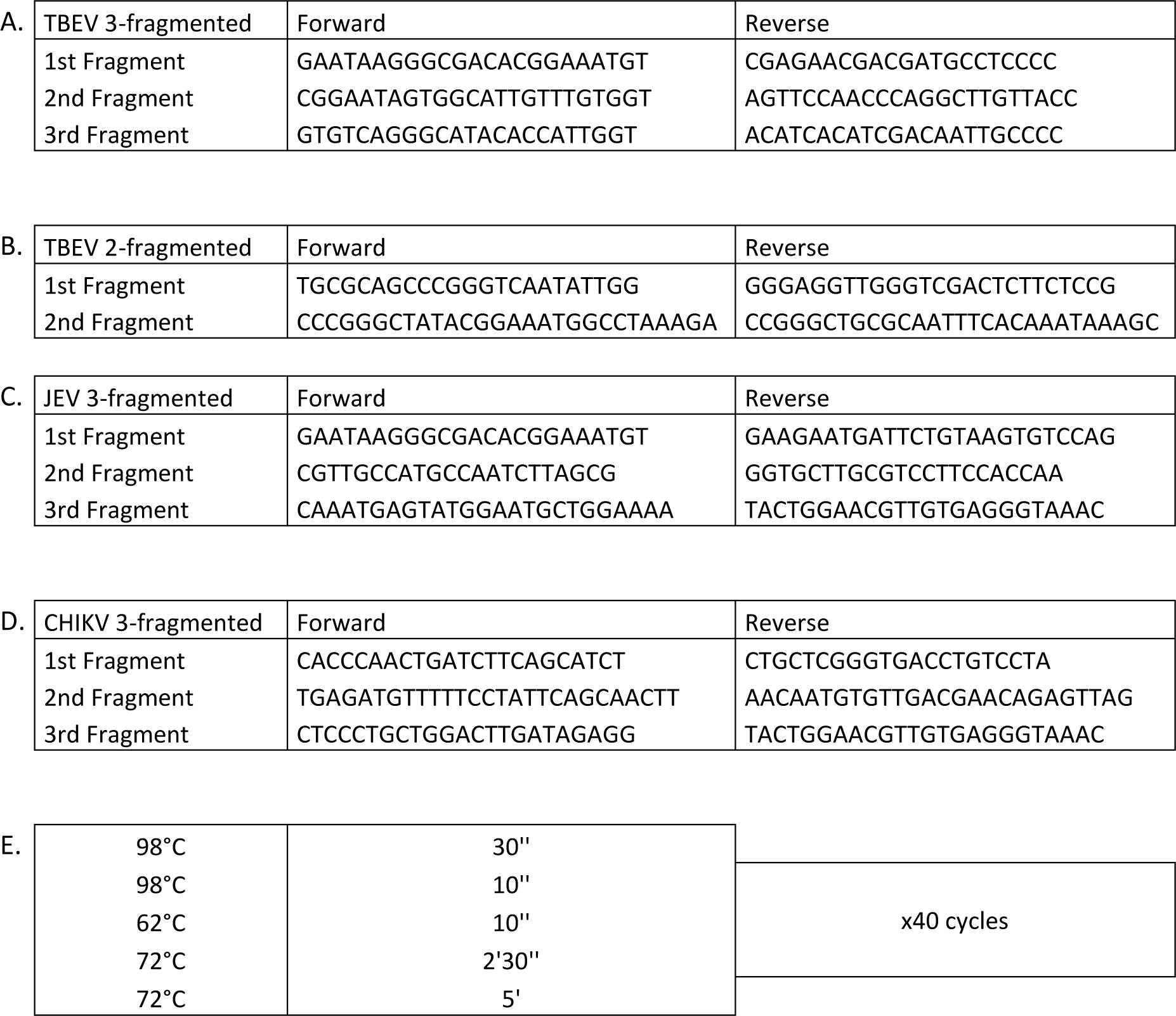
Primer pair sequences (A-D) and PCR cycle (E) to generate DNA fragments from reverse genetic systems.

**Table S2:**
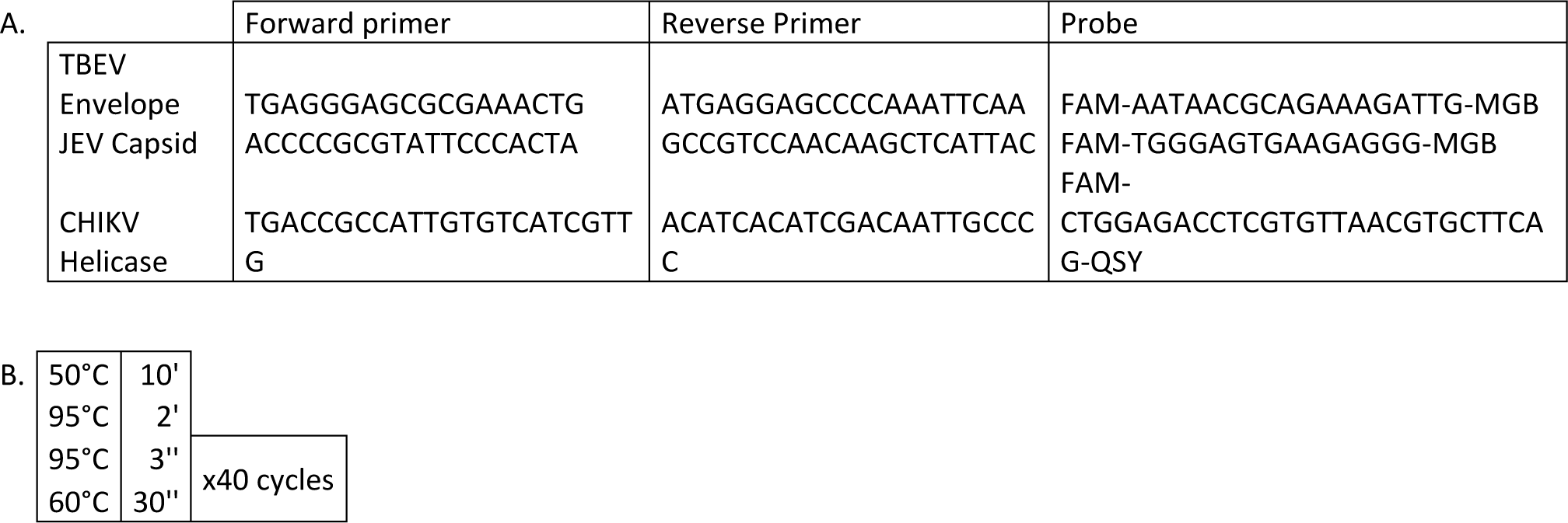
Viral detection systems (A) and cycle (B) for viral detection.

**Table S3:**
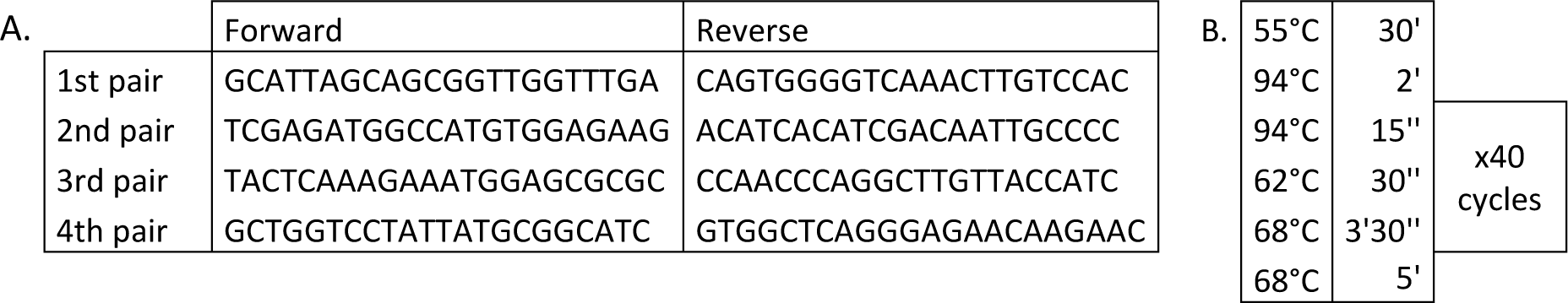
Primer pair sequences (A) and RT-PCR cycle (B) for next generation sequencing.

**Table S4:**
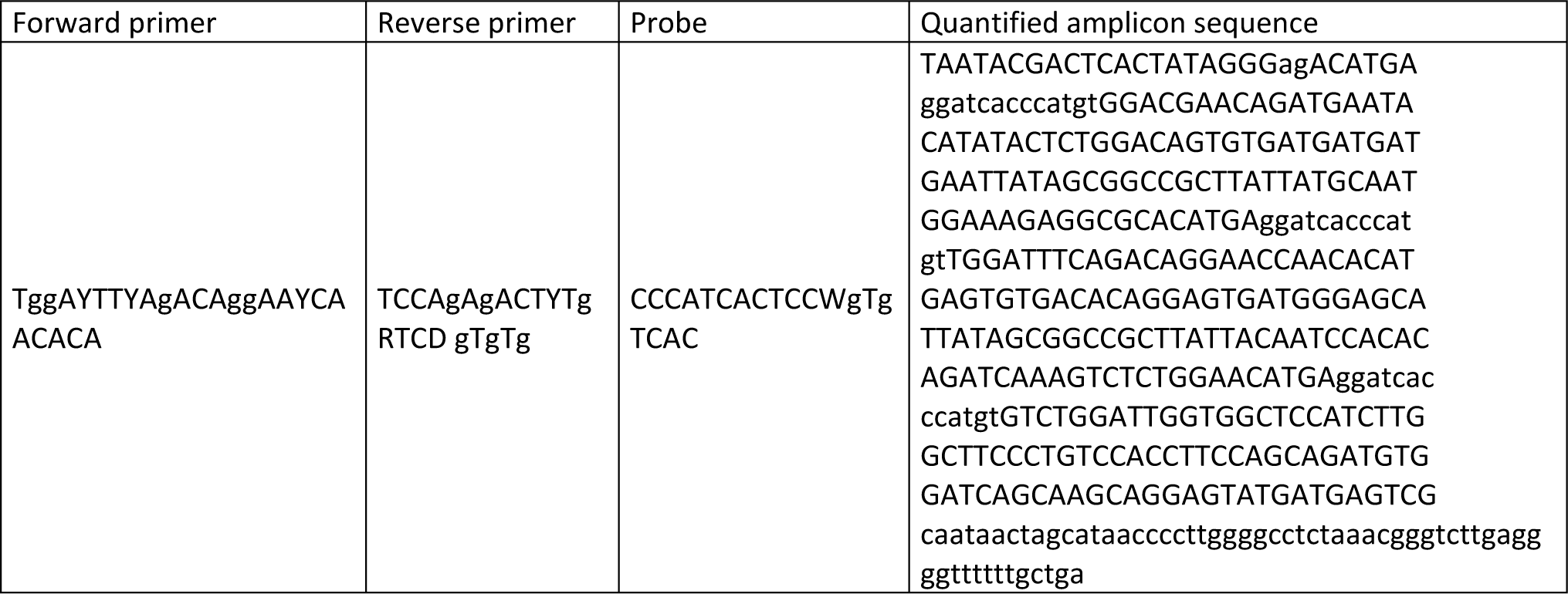
Quantification system used to quantify viral RNA yields for in vitro kinetic assay.

**Table S5:**
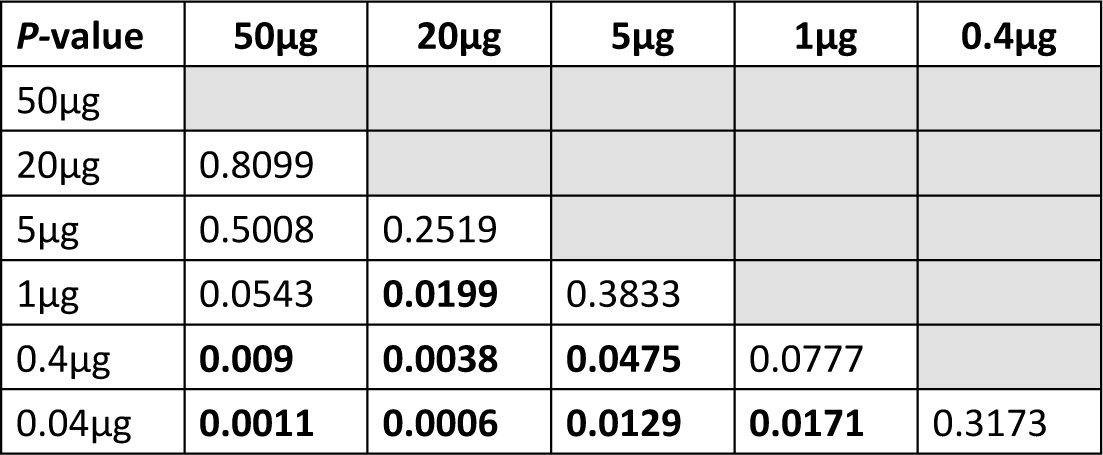
Statistical analysis (Log-rank test) of survival curves from groups presented in figure 1.

**Table S6:**
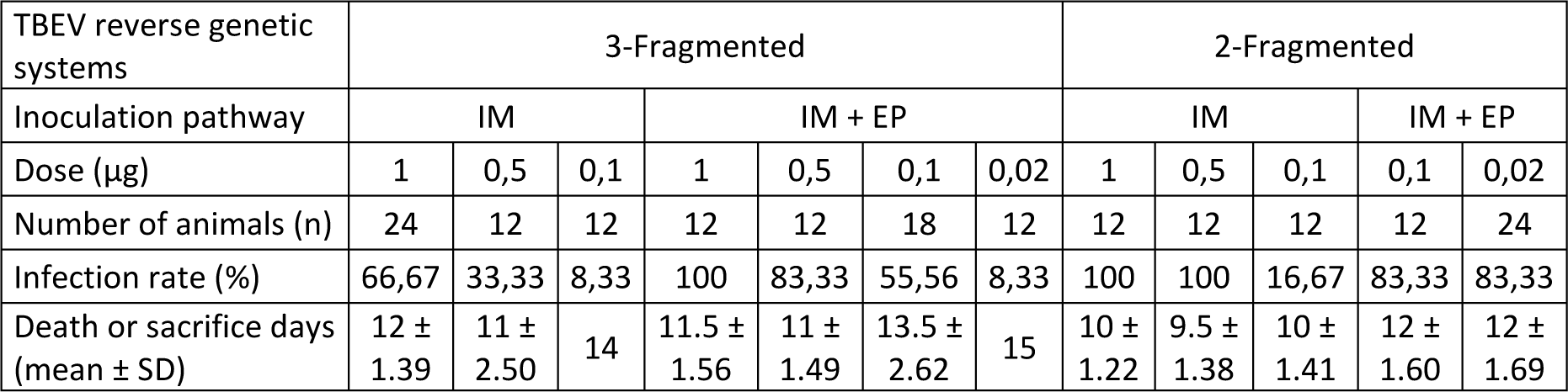
Survival data of animal experiments illustrated in figure 3 and Figure S7.

**Table S7:**
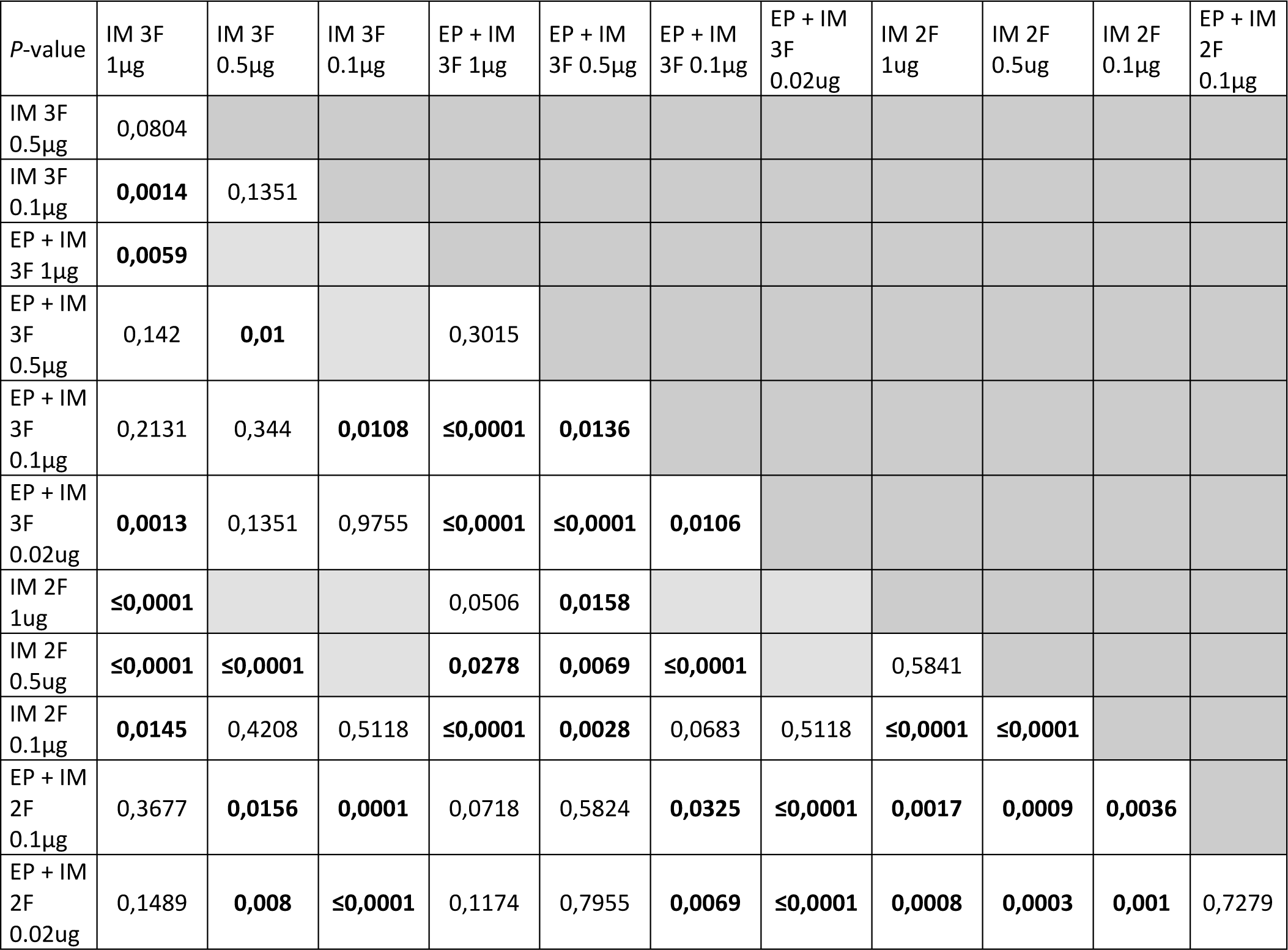
Statistical analysis (Log-rank test) of survival curves illustrated in Figure S7 and presented in figure 2.

**Table S8:**
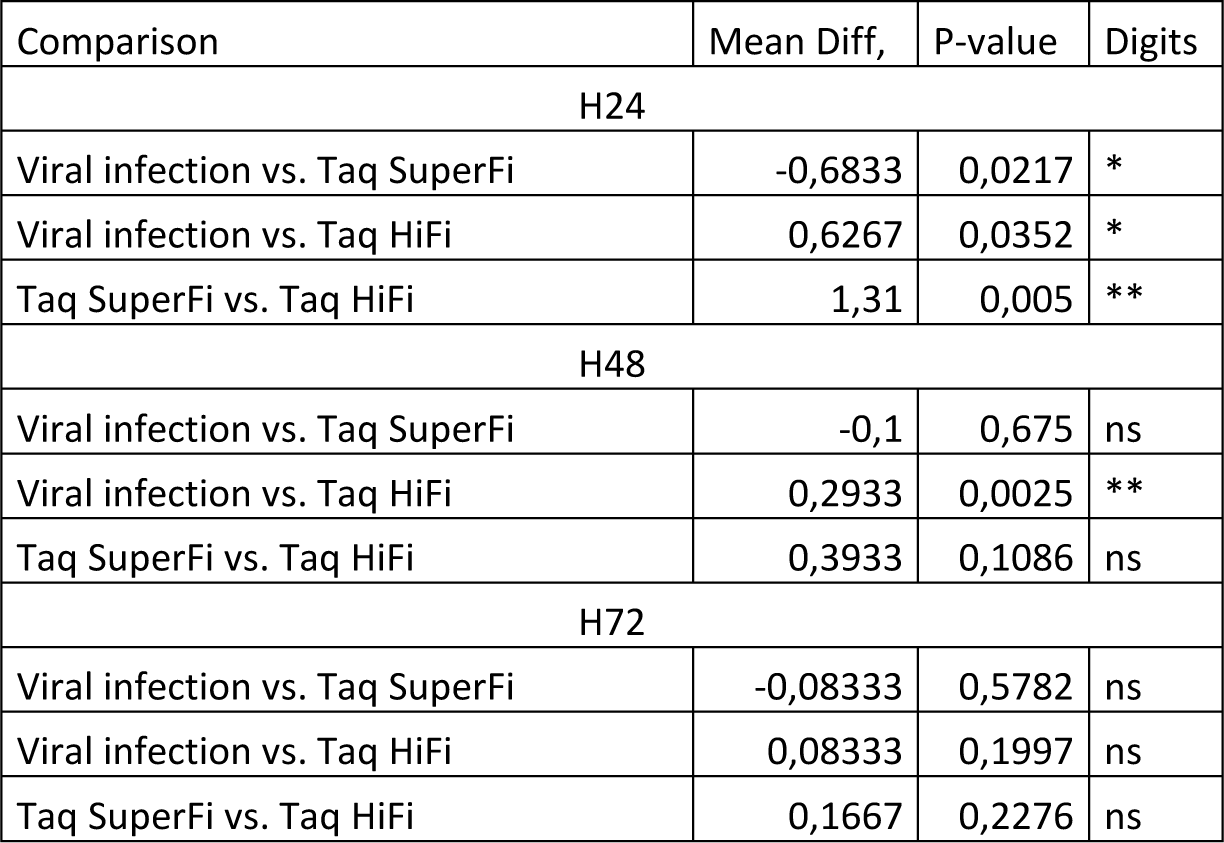
Statistical analysis (Two-way ANOVA, tukey’s multiple comparisons test) of in vitro replication fitness in Figure S4.

